# Protective human anti-poxvirus monoclonal antibodies are generated from rare memory B cells isolated by multicolor antigen tetramers

**DOI:** 10.1101/2022.06.04.494786

**Authors:** Xiuling Gu, Yufan Zhang, Wei Jiang, Dongfang Wang, Jiao Lu, Guanglei Gu, Cheng-Feng Qin, Min Fang

## Abstract

Smallpox, an epidemic disease caused by *Orthopoxvirus variola*, was eradicated worldwide through immunization. The immunization against smallpox was discontinued in 1980. However, incidences of monkeypox virus infection in humans have occurred sporadically, and there is also great fear that engineered forms of poxvirus could be used as biological weapons. Therefore, monoclonal antibodies against poxvirus are in urgent need for the detection and treatment of poxvirus infection. The vaccinia virus extracellular envelope protein A33 is a potential candidate for subunit vaccine. We used multi-fluorescence-labeled tetrameric A33 antigen to identify rare poxvirus specific memory B cells from PBMC of volunteers with vaccinia virus immunization more than 40 years ago. Despite extremely low frequencies of the poxvirus specific memory B cells, we successfully sorted A33 tetramer-labeled single memory B cells and reconstructed antibodies with single-cell RT-PCR of the B-cell receptor. Among the monoclonal antibodies, one clone H2 exhibited high specificity and affinity with A33. H2 efficiently inhibited viral infection and spread in cells. Passive immunotherapy of H2 in mice protected mice from lethal infection when administered either prophylactically or therapeutically. These results suggest the potential of the anti-A33 human antibody-based detection and therapeutics for poxvirus infection.

## Introduction

The *genus Orthopoxvirus* (OPV) is comprised of notable members such as variola virus (VARV), monkeypox virus (MPXV), vaccinia virus(VACV), cowpox virus (CPXV), and ectromelia virus (ECTV) [1]. Smallpox, caused by VARV, had periodically led to devastating epidemics throughout history and was eradicated following a worldwide vaccination campaign [2,3]. It has been more than four decades since the eradication of smallpox; vaccination programs have been halted and immunity has waned. Monkeypox is a zoonotic disease caused by MPXV [4]. In May 2022, multiple cases of MPXV infection had been reported in the United States and several European countries. Furthermore, there were some other zoonotic diseases including VACV in Brazil, CPXV in Europe, and buffalopox virus (BPVX) in India [5]. More seriously, the potential release of variola virus or engineered forms of poxvirus as an act of bioterrorism or the gradual transfer of zoonotic viruses from local hosts to humans emphasizes the importance of preparing for future poxvirus outbreaks.

VACV has been used as a live-attenuated vaccine to eradicate smallpox. Although VACV is an efficient vaccine, it routinely causes a pustular skin lesion, frequently induces lymphadenopathy and fever, and occasionally results in life-threatening disease [6]. Antibodies have been used as therapeutic drugs for more than 100 years. Monoclonal antibodies (mAbs) are effective biologics for use as therapeutic and diagnostic reagents [7]. Murine mAbs have a short half-life when used in humans and are relatively poor recruiters of effector function, such as antibody-dependent cellular cytotoxicity, and complement-dependent cytotoxicity [8]. Chimeric mAbs are constructed by coupling the rodent antigen-binding variable (V) domains to human constant (C) domains, are still 30% rodent in sequence and can elicit a significant anti-globulin response [4,9]. Humanized mAbs are created by transplanting the complementarity determining regions (CDRs), which are the hypervariable loops involved in antigen binding, from rodent antibodies into human V domains. However, the process to generate these molecules is arduous and has limitations [10]. With the development of new technology, it becomes feasible to directly clone human antibody from a single B cell. Human mAbs have a longer circulation time in comparison with murine or chimeric antibodies [11].

There are two forms of infectious OPV: extracellular enveloped virion (EV) and intracellular mature virion (MV) [12]. MV particles are responsible for host-to-host spread, whereas EV particles are important for virus dissemination within the host as well as in cultured cells [6,13,14]. The reported target antigens recognized by neutralizing antibodies include A33 and B5 on the surface of EV; L1, H3, A27, D8, A28, B7, and A17 of MV [15]. Among them, A33 plays an important role in the spread of the virus between cells. Immunization with A33 in mice elicits antibody responses and protects susceptible mice from lethal ECTV infection [16].

Over the past two decades, single B cell antibody technologies have become an effective approach to sample naïve and antigen-experienced antibody repertoires generated in vivo [17]. Single B cell sorting and cloning of V_H_ and V_L_ is a powerful technology for generating neutralizing mAbs against infectious diseases, for example, mAbs against H1N1 or H5N1 influenza virus, SARS, and SARS-COV-2 coronaviruses [18-21]. Normally, fluorescently labeled antigen has been used to identify B cells with particular BCR specificities by flow cytometry. However, the frequency of most specific memory B cells is low and signals generated by fluorescently labeled antigens are typically not bright and tend to overlap with unlabeled cell populations. Franz B et al generated fluorescent antigen tetramers to isolate low frequency memory B cells [22].

Here, we used multiple fluorescent A33 tetramers to identify and isolate rare memory B cells from PBMC of a vaccinated volunteer.

## Materials and Methods

### Ethics Statement

This study for human peripheral blood was performed with the informed consent of the participant. The experimental design and protocols used for human peripheral blood in this study were approved by the Research Ethics Committee of the Institute of Microbiology, Chinese Academy of Sciences (permit number APIMCAS2019056). The animal protocols used in this study were approved by the Research Ethics Committee of the Institute of Microbiology, Chinese Academy of Sciences (permit number APIMCAS2017034).

### Cells and viruses

293T cells (ATCC, CRL-3216) and BSC-1 cells (ATCC, 3168) were cultured in complete DMEM medium. 293-F cells (Invitrogen, R790-07) were cultured in SMM 293-TI. All cells were cultured at 37°C in 5% CO_2_.

Initial stocks of ECTV Moscow and VACV Western Reserve were obtained from Dr. Luis Sigal (Thomas Jefferson University) and amplified and quantified as described [23].

### Production and Biotinylation of A33

The coding sequences for the extraviral domain of VACV Western Reserve A33 [24] were amplified by PCR from genomic DNA. For A33 the forward and reverse primers used were 5’AAACCATGGGCCATCACCATQCACCATCACTGCATGTCT-GCTAACGAGGCTG3’ and 5’AAAGGATCCGTTCATTGTTTTAACACAAAAATA CTTTC3’, respectively. Avi-tag (GLNDIFEAQKIEWHE)[25] was fused to the C-terminus of A33 to mediate biotinylation. The expression, purification and renaturation of A33 were performed as previously described [16]. Biotinylation of A33 protein was performed by using the biotin-protein ligase kit (GeneCopoeia™, B1001) according to the manufacturer’s instructions.

The coding sequences for the extraviral domain of VARV A33 with N terminus 6×His tag and C-terminus Avi-tag were synthesized by GenScript and directly cloned into the PET-28a(+). The purification and biotinylation of VARV A33 were the same as that of VACV A33.

### Mice and infection

Female C57BL/6 (B6) and BALB/c mice (6-8 weeks) were purchased from Vital River, China. For infection experiments, mice were transferred to a biosafety level 2 room. B6 mice were infected in the left footpad with 30μl of PBS containing 3000 plaque forming unit (PFU) of ECTV, or infected intraperitoneal (i.p.) with 5×10^6^ PFU VACV.

Female BALB/c mice were i.p. infected with 5×10^6^ PFU VACV. H2 IgG was administered at 22 mg/kg into the indicated mice.

### Production of antisera

Anti-virus antisera were produced by infection B6 mice with ECTV or VACV. The sera were collected at five weeks pi, and stored in aliquots at -80°C.

### Flow cytometry and tetramer preparation

All the tetramers were prepared freshly for each experiment. Biotinylated A33 was incubated with APC-conjugated streptavidin (SA-APC), APC/Cy7-conjugated streptavidin (SA-APC/Cy7), or pacific blue-conjugated streptavidin (SA-PB, eBioscience) for 3 min at room temperature at a molar ration of 4:1, respectively. The fluorescent A33 tetramers were then used together with other antibodies for cell staining. The splenocytes of mice were stained as described previously [23].

### Single-cell sorting

Blood samples were donated by a healthy volunteer. PBMC were isolated and stained with the indicated cell surface markers. Single memory B cells were sorted into 96-well PCR plates as described previously [25].

### Single-cell RT-PCR and amplification of antibody variable region sequences

The genes encoding IgG V_H_ and V_L_ chains were amplified by RT and nested PCR using the previously reported method [25]. All PCR products were purified and sequenced. Sequences were analyzed using the VBASE2 (http://www.vbase2.org) to identify variable region gene segments.

### Constructing and expression of single chain antibody (scFv)

The scFvs were constructed by connecting paired V_L_ and V_H_ with a (GGGS)_2_ linker, an Ig leader sequence (METDTLLLWVLLLWVPGSTGD) was used as a signal peptide. A flag tag was fused to the C-terminus of scFv for easy detection. The constructs were cloned into pcDNA3.1 vector. The plasmids were transfected into 293F cells, and the cell supernatants were collected after 48-72 hours.

### Production of H2 IgG

H2 IgG was generated by combining V_H_ and V_L_ with constant regions of IgG1 heavy and light chain, respectively. The constructed heavy and light chain plasmids were co-transfected into 293F cells. H2 IgG was purified by protein A according to the manufacturer’s recommendations.

### ELISA

ELISA plates were coated with VACV A33, VARV A33 (50 μg/ml), ECTV (1×10^7^ PFU/ml), VACV (1×10^7^ PFU/ml), PR8 (1×10^7^ PFU/ml) or BSA (50 μg/ml) as indicated. ELISA was performed as described previously [16].

### Comet inhibition assay

BSC-1 cells were infected with ECTV (MOI=0.01), the indicated dilutions of antisera or antibodies were added. The assay was performed as described previously [16].

### Surface plasmon resonance analysis

The affinity between H2 and A33 was measured at room temperature using a Biacore T100 system with CM5 chips (GE Healthcare) according to the manufacturer’s recommendations.

### Histopathology

Histopathology was performed as described previously [26].

### Statistics

Statistical analysis was performed using Prism software (GraphPad). All statistical analyses were performed using an unpaired two-tailed Student’s t-test or two-way ANOVA test as applicable.

## Results

### Production and biotinylation of VACV A33

Previous studies demonstrated that immunization of A33 protein protects susceptible mice from lethal mousepox [16]. Importantly, the A33 extraviral region of VARV, VACV, MPXV, and ECTV is highly homology (92.68%) (Fig. 1A). The coding sequence of the extraviral domains of VACV A33 with a His-tag at the N terminus and an avi-tag at the C terminus was cloned into the expression vector pET-28a (+) (Fig. 1B) for producing recombinant A33 in *E*.*coli*. The avi-tag mediates biotinylation by BirA enzyme. A33 was successfully expressed and purified (Fig.1C and 1D). Next, A33 was biotinylated and verified by Western blot (Fig. 1E). Biotinylated A33 was recognized by both anti-ECTV and anti-VACV antisera (Fig.1F). These results suggest that biotinylation does not affect the binding of A33 to anti-OPV antibodies.

**Figure 1.**
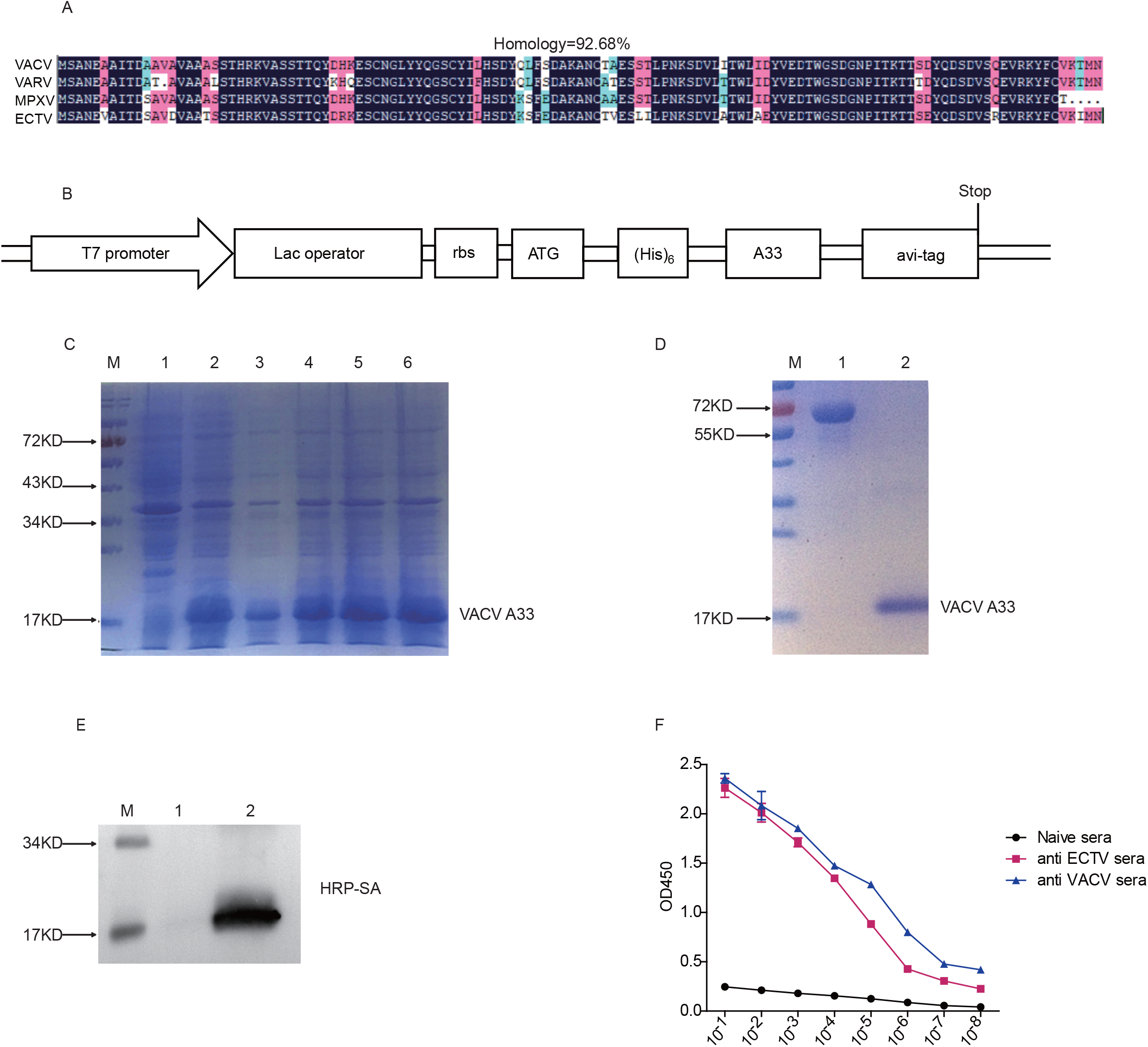
Production and biotinylation of A33. (A) DNAMAN software comparison of the homology of amino acid sequences of A33 extracellular region from VARV, VACV, MPXV, and ECTV. (B) Scheme of the expression vector. (C) Expression of A33 in BL21(DE3) cells induced with IPTG. Lane 1: Control BL21(DE3) cells (vector). Lane 2, 3, 4, 5, 6: different clones of BL21(DE3) cells transfected with pET/A33. (D) Purification of A33. Lane 1: 1mg/ ml BSA for comparison; Lane 2: Purified and refolded A33 protein. (E) Western Blot detection of A33 biotinylation. Lane 1: A33 protein; Lane 2: Biotinylated A33 protein. (F) biotinylated A33 was recognized by anti-ECTV sera and anti-VACV sera. Sera from naive mice were used as a negative control.

### Tetramer-based screenings of A33-specific memory B cells

To improve the efficacy and specificity of the screening for rare A33 specific memory B cells, we generated multiple colors labeled A33 tetramers. Biotinylated A33 was incubated separately with different fluorescent conjugated streptavidin. The generated A33 tetramers were then used together with a panel of mAbs to identify the A33 specific memory B cells (Fig. S1A). To verify the feasibility of this strategy, we first infected B6 mice with 3000 pfu ECTV in the footpad. 4-5 weeks post infection, leukocytes in the spleens were isolated for flow cytometry analysis. As shown in Fig. S1B, we first gated on the CD3^-^B220^+^CD27^+^ B cells, the A33-specific B cells were selected as the A33-Tet-PB^+^A33-Tet-APC^+^ double positive cells. In the naive B6 mice, the proportion of A33-specific B cells was 0.53%. Surprisingly, the proportion of A33-specific B cells reached 39.7% in the ECTV infected mice. The double staining of A33 tetramers defined a concentrated and clear cell population. This result was striking and the importance of anti-A33 antibody responses during ECTV infection was emphasized. We further detected A33-specific memory B cells after VACV infection. B6 mice were i.p. infected with 5×10^6^ PFU VACV. 4-5 weeks post infection, leukocytes in the spleens were isolated for flow cytometry analysis. We gated on B220^+^IgM^-^CD27^+^ class switched memory B cells. While in the VACV infected mice, A33 specific memory B cells comprised around 16% of total memory B cells (Fig.S1C). Therefore, anti-A33 antibody responses account for a high proportion of the whole anti-poxvirus antibody responses.

Next, we used multiply color A33 tetramers to isolate memory B cells from the PBMC of a healthy volunteer who was vaccinated against smallpox more than 40 years ago. As shown in Fig. S1D, A33-Tet-APC staining showed around 1% of A33 specific memory B cells among the total CD19^+^CD27^+^IgM^-^ memory B population, however, the staining was not very bright and contained some background staining. A33-Tet-PB staining showed a similar pattern. Whereas A33-Tet-PB and A33-Tet-APC double staining defined a very rare cell proportion (0.045%) in the diagonal, which sharply reduced background staining. Single A33-Tet-PB^+^A33-Tet-APC^+^ double positive memory B cells were sorted by flow cytometry and stored in 96-well plates for subsequent PCR amplification.

### Amplification of antibody variable region and construction of single-chain antibody

Single B cell mRNA amplification and nested PCR to amplify Ig V_H_ and V_L_ genes were performed as described previously [25]. From a total of 96 single cells, 19 pairs of heavy chain and light chain variable fragments were amplified, corresponding to an overall PCR efficiency of around 20% (Fig. S2A). We further analyzed the V, D, and J regions of 19 mAbs by the IMGT database. The results showed that all 19 antibody light chains were κ, and IGHV3-7*05 accounted for the highest proportion in the V region of IGH, IGHJ5*01 in the J region was the highest, and the proportions of IGKV1-5*03 and IGKJ2*01 were higher in IGK (Fig. S2B).

The CDRs of V_H_ and V_L_ domains determine the antigen specificity of the Ig [27,28]. We then compared the sequences of the amplified 19 pairs of V_H_ and V_L_ (Fig. S2C). A7, A8, A10 and A11 had identical V_H_ and V_L_ indicating that the cells might come from one original B cell clone. B9 and H2 also had identical V_H_ and V_L_. In addition, nine clones (B1, B2, B7, B11, C1, C3, C4, C6, and D7) differed from A7 in 1-8 amino acids in the framework and CDR regions. The remaining B3, C2, and C8 demonstrated a larger number of amino acid differences, indicating distant clonal relationship. Thus, based on the difference in sequences, especially within CDRs, we selected 9 pairs of V_H_ and V_L_ to construct a single-chain antibody (scFv) (Fig.S2D) for rapid and easy screening of the antigen specificity of the Ig.

We transfected the 9 scFv plasmids into 293F cells, respectively. Among the 9 constructs, only H2, A7, B3, and B6 were expressed (Fig. S2E). Next, ELISA plates were coated with A33 protein and tested for recognition by the scFvs. As shown in Fig.2A (left panel), only H2 and A7 bound to A33, but the affinity of A7 was lower than that of H2. 293F cell culture supernatant was used as negative control. As H2 showed higher binding capacity to A33, we next determined the binding specificity of H2 scFv. ELISA plates were coated with A33, influenza virus A/Puerto Rico/8/34 (PR8), or vesicular stomatitis virus (VSV). H2 is only bound to A33, not PR8 or VSV (Fig.2A, right panel). A33 is highly conserved between OPVs. As expected, H2 scFv bound both ECTV and VACV (Fig. 2B).

**Figure 2.**
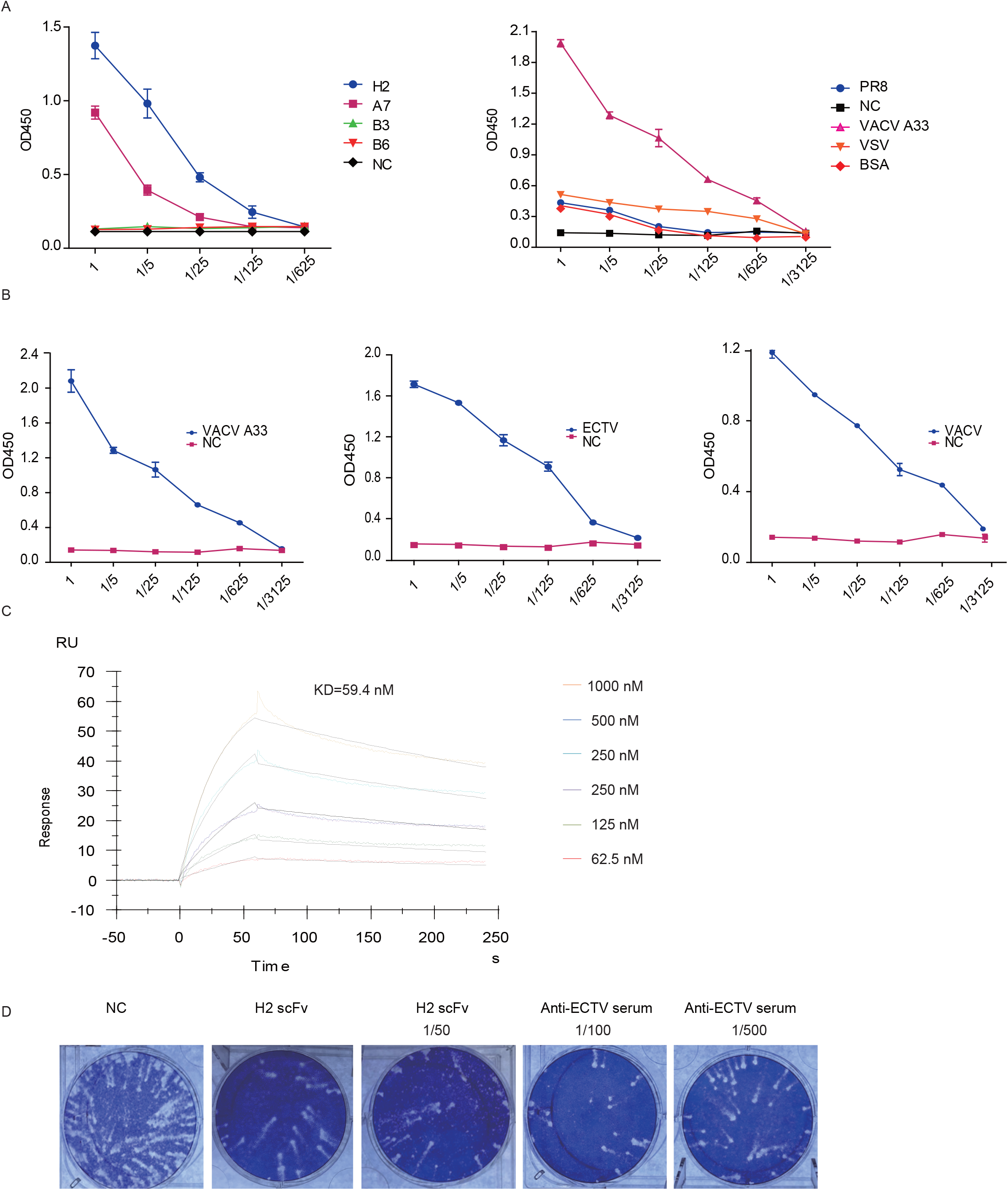
Functional identification of ScFvs. (A) ELISA plates were coated with A33 and then incubated with the indicated scFvs (left panel); ELISA plates were coated with A33, PR8, VSV, or BSA and then incubated with H2 scFv (right panel). (B) ELISA plates were coated with A33, ECTV, or VACV, and then incubated with serial dilutions of H2 scFv. Uncoated wells were used as negative control (NC). (C) Representative SPR sensorgrams of H2 scFv binding to A33. (D) Comet inhibition assay.

### H2 scFv blocks virus replication in cell culture

We used surface plasmon resonance (SPR) to determine the affinity of H2 scFv to A33. As shown in Fig. 2C, the binding affinity (KD) between H2 scFv and A33 was 59.4 nM. Our previous studies demonstrated that antisera against A33 inhibit comet formation in ECTV infected cells [16]. Therefore, we determined whether comet formation by ECTV could be inhibited by H2 scFv. As shown in Fig. 2D, H2 scFv inhibited comet formation as well as viral infection. The anti-ECTV antisera were used as a positive control. Anti-ECTV antisera or H2 scFv both resulted in markedly reduction of plaques in addition to the inhibition of comet. It may be because the antisera or scFv block released virions to infect adjacent cells, thus preventing the formation of plaques. Previous studies have shown that Abs against the EV B5 protein are mainly responsible for the EV neutralizing capacity of VACV as measured by a plaque reduction assay [29]. Our experiments demonstrated that Ab against EV A33 also can reduce plaque formation. Collectively, H2 scFV shows functional activity in vitro.

### Purification and functional characteristics of H2 IgG

Next, we generated H2 IgG by combining V_H_ and V_L_ with constant regions of IgG1 heavy and light chains, respectively. The purified H2 IgG was verified by SDS-PAGE (Fig. 3A). H2 IgG specifically recognized the A33 protein, ECTV, and VACV, but not PR8 or VSV, which was consistent with the results of scFv (Fig. 3B and 3C). In addition, H2 IgG inhibited comet formation and also viral replication in ECTV-infected BSC-1 cells (Fig. 3D). Next, we wanted to verify whether H2 IgG can bind to VARV A33. The coding sequences for the extraviral domain of VARV A33 were synthesized. VARV A33 was expressed and purified (Fig. 5E and 5F). As shown in Fig. 3G, H2 IgG recognized the VARV A33. This result suggests that H2 IgG may inhibit the spread of the VARV as well and is a broad protective mAb against OPV.

**Figure 3.**
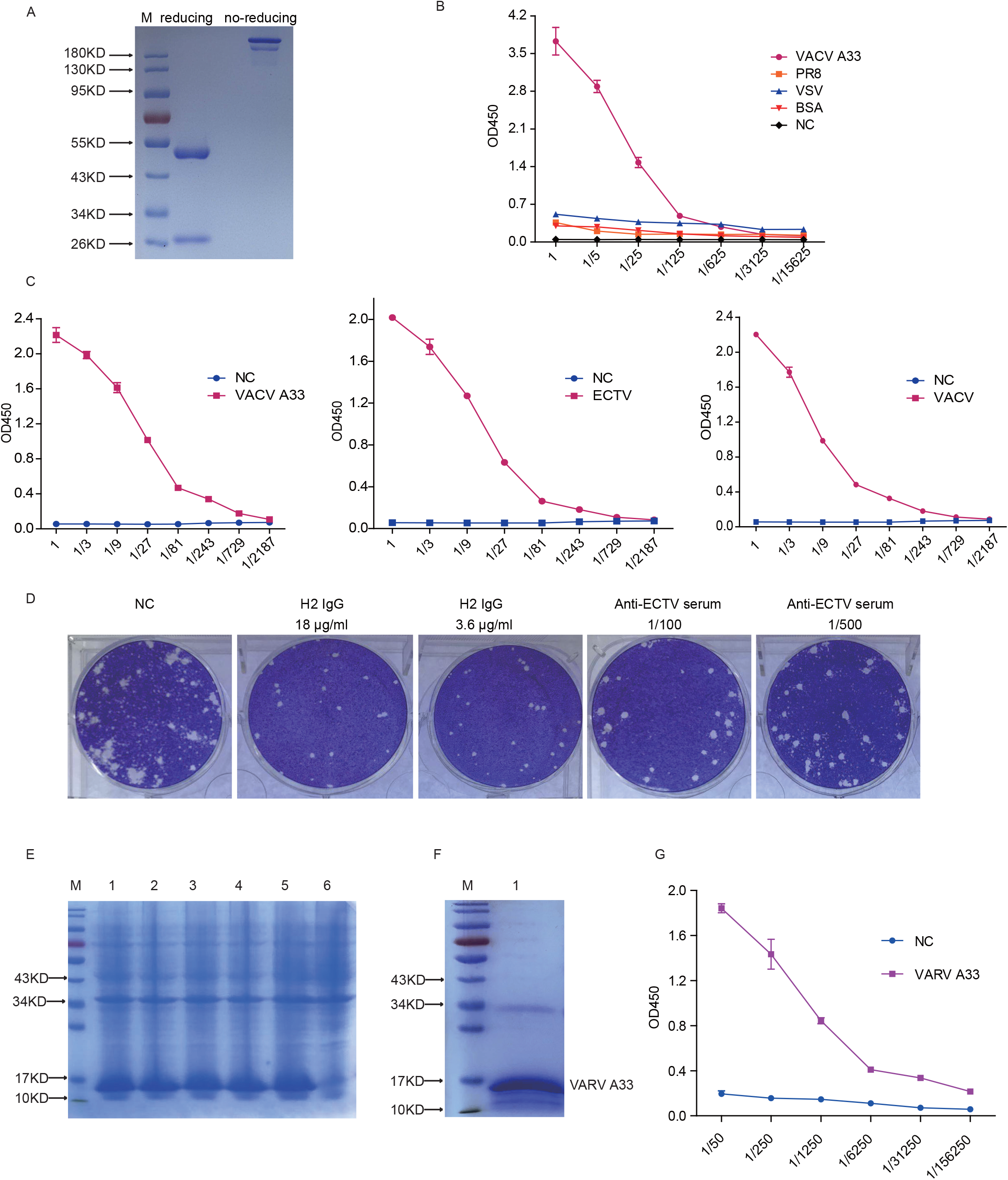
Purification and functional identification of H2 IgG. (A) SDS-PAGE of the purified H2 IgG under reducing (left) and non-reducing conditions (right). (B) The wells of ELISA plates were respectively coated with A33, VSV, PR8, or BSA and then incubated with serial dilutions of H2 IgG. (C) The wells of ELISA plates were coated with A33, ECTV, or VACV, respectively, and then incubated with serial dilutions of H2 IgG. (D) Comet inhibition assay. (E) Expression of VARV A33 in BL21 (DE3) cells induced with IPTG. Lane 1, 2, 3, 4, 5: different clones of BL21 (DE3) cells transfected with pET/VARV A33. Lane 6: Control BL21 (DE3) cells (vector). (F) Purification of VARV A33. Lane 1: Purified and refolded VARV A33 protein. (G) The wells of ELISA plates were coated with VARV A33 and then incubated with serial dilutions of H2 IgG.

### H2 IgG protects mice against VACV infection

To test whether H2 IgG can protect mice from OPV infection, BALB/c mice were i.p. infected with 5×10^6^ PFU VACV. H2 IgG or PBS was administered 4 hours before and 48 hours after infection (Fig. 4A). As shown in Fig. 4B, both groups of mice lost weight after infection, the weight loss in the H2 IgG group remained relatively stable from 1 dpi to 5 dpi, and started to recover the weight from day 6. However, the mice of the PBS group continued to lose weight from 1 dpi to 10 dpi. Only around 20% of the mice in the PBS-treated control group survived after infection, while the mice injected with H2 IgG had a survival rate of up to 80% (Fig. 4C). Thus, H2 IgG significantly protected mice from VACV infection when administered prophylactically.

**Figure 4.**
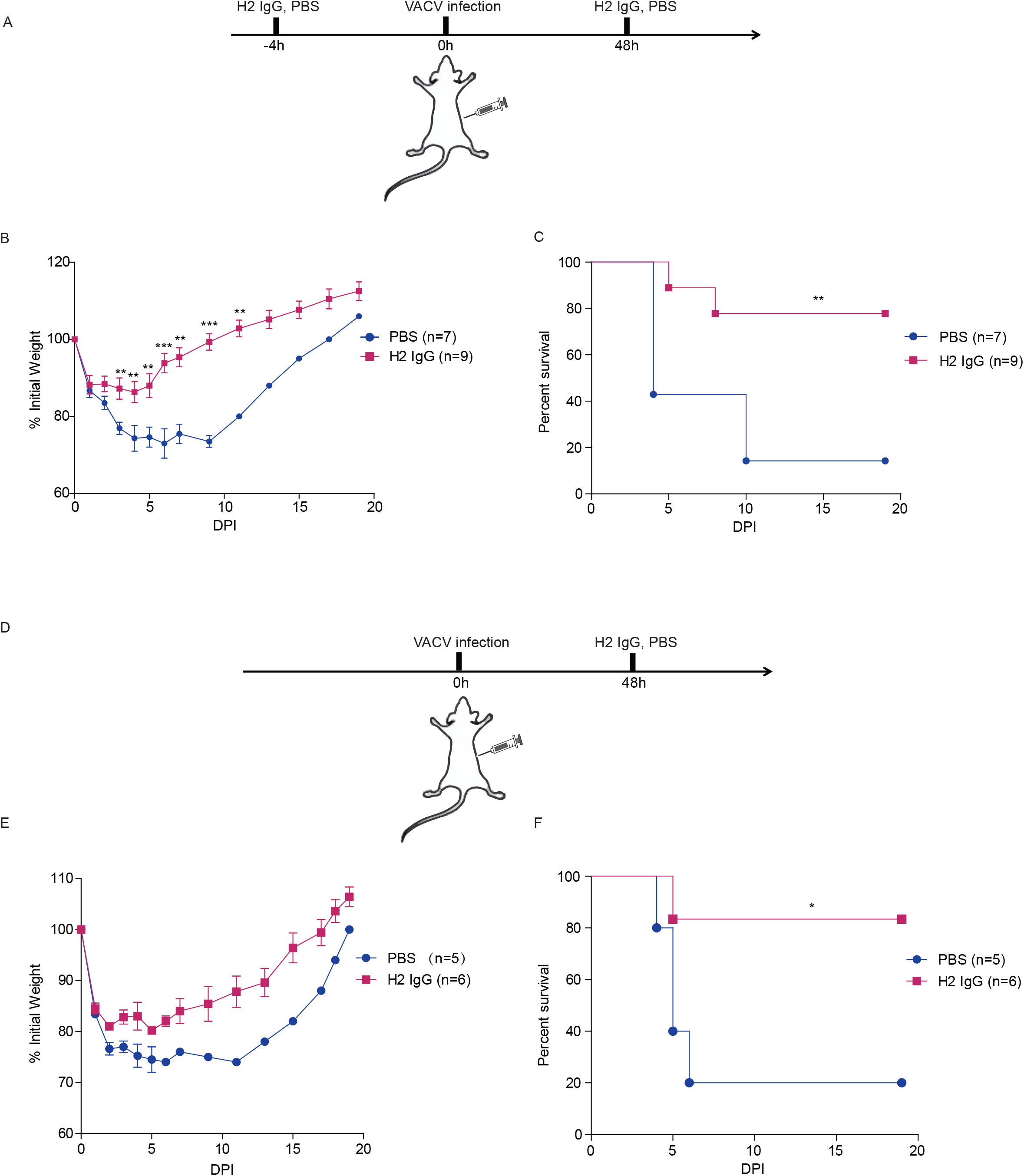
Prophylactic and therapeutic protection in mice by H2 IgG. Female BALB/c mice were i.p. infected with 5×10^6^ PFU VACV. H2 IgG was administered in mice (22 mg/kg) at the indicated time points. (A) Schematic diagram of prophylactic treatment with H2 IgG in mice. (B) Weight change following VACV infection. (C) Survival curves following VACV infection. (D) Schematic diagram of treatment with H2 IgG in mice. (E) Weight change following VACV infection. (F) Survival curves following VACV infection. Data are from at least two independent experiments with 3-5 mice per group in each experiment. *p<0.05, **p<0.01, ***p<0.001.

Next, we tested whether H2 IgG can protect mice after VACV infection. BALB/c mice were infected with VACV. After 48h, the mice were injected with H2 IgG or PBS (Fig. 4D). Again, both groups of mice experienced weight loss after infection, however, the H2 IgG group started to regain the weight from 5 dpi (Fig.4E). Again, H2 IgG administration after VACV infection significantly protected mice from death (Fig. 4F). Collectively, these results demonstrated that H2 IgG can protect mice from VACV infection both prophylactically and therapeutically.

### H2 IgG promotes anti-VACV T cell and Ab responses

T cell responses are known to be important for protection against OPV infections [23,30,31]. Therefore, we next determined whether administration with H2 IgG has any effects on the T cell responses. Mice were first administrated with H2 IgG, or PBS, then challenged with VACV and 7 days later, their spleens were collected. Compared to naïve mice, the spleens of the mice in the PBS group were reduced, while the spleen of the mice in the H2 IgG group was significantly enlarged. The total number of splenocytes in naive mice was 3.9±0.15×10^7^, the PBS group was 2.6±0.35×10^7^, the H2 IgG group was 8.3±0.29×10^7^, which increased significantly (Fig. 5A). Moreover, after VACV infection, the percentage of CD4^+^ T cells decreased. However, the CD4^+^ T cells in H2 IgG group increased, while the PBS group decreased. In addition, the percentage of virus-specific CD4^+^ T cells increased in H2 IgG group compared with that of PBS group as determined by intracellular IFN-γ staining following in vitro restimulation with VACV-infected cells (Fig. 5B).

**Figure 5.**
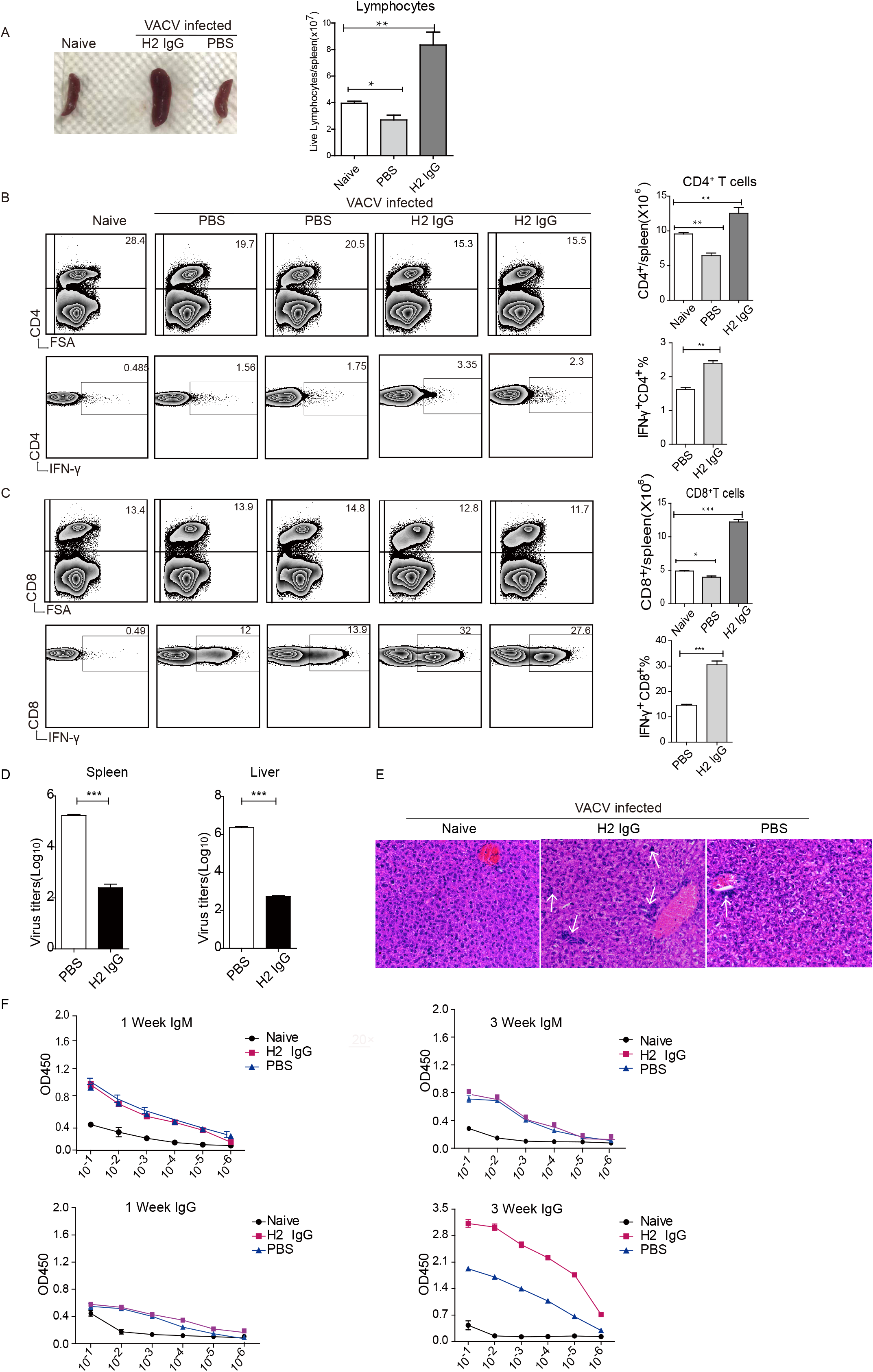
H2 IgG promotes anti-VACV T cell and Ab responses. (A) Indicated mice were administered with H2 IgG or PBS before VACV infection. The mice were euthanized at 7 dpi and the size of the spleens was observed (left panel). Total lymphocytes per spleen were counted (right panel). (B) Left panel: Flow cytometry analysis showing the proportion of CD4^+^ T cells in the spleen, and the proportion of IFN-γ^+^ cells among CD4^+^ T cells. FSA: forward scatter amplitude. Stained as indicated. Right panel: Total CD4^+^ T cells per spleen and proportion of IFN-γ^+^ cells among CD4^+^ T cells. (C) Left panel: Flow cytometry analysis showing the proportion of CD8^+^ T cells in the spleen, and the proportion of IFN-γ^+^ cells among CD8^+^ T cells. Stained as indicated. Right panel: Total CD8^+^ T cells per spleen and proportion of IFN-γ^+^ cells among CD8^+^ T cells. Numbers indicate the proportion in the nearest quadrant. (D) Virus titers in spleens and livers were determined on 7 dpi. (E) H&E stains of the liver (original magnification,×20) on 7 dpi. The arrow pointed to the lymphocyte infiltration area. (F) anti-VACV IgM or IgG Ab responses were determined at 1 week and 3 weeks p.i. Data are from at least two independent experiments with 3 mice per group in each experiment. *p<0.05, **p<0.01, ***p<0.001.

The percentages of CD8^+^ T cells remained relatively stable after VACV infection, while the cell number of CD8^+^ T cells in H2 IgG group increased. Markedly, the percentage of IFN-γ^+^ CD8^+^ T cells increased in H2 IgG group, reaching around 30%, which was significantly higher than that of the PBS group (Fig. 5C). Therefore, administration of H2 IgG prevented lymphocyte depopulation and promoted stronger anti-VACV CD4^+^ and CD8^+^ T cell responses.

Next, we determined the viral loads in spleen and liver. As shown in Fig. 5D, H2 IgG administration resulted in significantly decreased viral titers in both spleen and liver. Moreover, even though both groups had necrotic foci in their livers, those of the H2 IgG group were more heavily infiltrated with lymphocytes than that of the PBS group (Fig. 5E). In addition, at 1 week pi, anti-VACV IgM was detected in both infected groups, while almost no anti-VACV IgG was detected as we only detected murine IgG. However, the IgG level in the H2 IgG group was substantially higher than that of the PBS group at 3 weeks pi (Fig. 5F). Therefore, H2 IgG promoted anti-VACV Ab responses as well. Together, these results demonstrated that administration of H2 IgG resulted in stronger T cell and Ab responses.

## Discussion

Previous studies have suggested that antibodies are sufficient to protect against OPV infections in mice and monkeys [32,33]. Vaccinia immune globulin (VIG) is manufactured from vaccinia vaccine-boosted plasma; however, this production method is not ideal because of its limited availability, and risk of contamination with blood-borne infectious agents [34,35]. Abs in VIG recognize many antigen targets, including surface proteins of both MV (A27, L1, H3, D8, A28, A13, and A17) and EV (B5 and A33) virion forms of VACV [36]. Protective mAbs had been reported against A33, B5, L1 and H3. Such as anti-B5 chimpanzee/human mAbs 8AH8AL and 8AH7AL [37], anti-L1 mouse mAb 7D11 [38]. Human mAbs against H3 and B5 were produced by using KM mice [39,40]. A large panel of OPV-specific human Abs was generated by hybridoma cell lines, the mAb panel contained Abs to at least 12 antigens: D8, B5, A33, H3, L1, A27, I1, A25, F9, A28, A21, and H5 [41]. Recently, two human scFvs (SC34 and SC212) that bound to VACV were isolated by a phage library constructed from the B cells of VACV vaccine-boosted volunteers [34].

EV membrane protein A33 plays an important role in effective cell to cell spread within the host [42,43]. A33 is also required for the proper formation of infectious EEV[43,44]. Anti-A33 Abs can be protective in vivo [15,16,45]. Anti-A33 chimpanzee/human mAbs 6C, 12C, and 12F exhibited higher protective efficacy than did a mouse anti-A33 mAb or human VIG [15]. Furthermore, 6C had been humanized [46]. Here, we sorted A33-specific memory B cells by using multicolor tetramer staining and got one mAb H2 with high specificity and affinity to A33.

The mAb H2 showed functional activity in vitro. Moreover, mAb H2 protected BALB/c mice from lethal VACV infection either administered prophylactically or therapeutically. T cell responses are known to be important in protection against OPV infections [47]. We found that the number of CD4^+^ and CD8^+^ T cells and also the T cell responses all increased in the mice administered with mAb H2. Further, administration of H2 also stimulated stronger Ab responses in the mice.

Another important observation of our study is the direct visualization of the anti-A33 Ab responses during OPV infections. The proportion of A33-specific B cells reached 39.7% in the ECTV infected mice. During VACV infection, A33 specific memory B cells comprised around 16% of total memory B cells. This demonstrated that anti-A33 antibody responses account for a high proportion of the whole anti-poxvirus antibody responses. As ECTV is a natural mouse pathogen, spreads massively in mice, the high proportion of anti-A33 Ab responses highlights the importance of anti-A33 Abs in protection. Since VARV specifically infects humans, and human being is the only natural host for VARV, anti-A33 Ab responses might also play an important role in curbing virus spread in vivo. As the A33 protein is highly conserved among OPVs, our H2 mAb might be used for treatment of complications of smallpox vaccination and a new approach to the prevention and treatment of OPV infections, including MPXV.

## Acknowledgments

We thank Yuanyuan Chen, Zhenwei Yang, and Bingxue Zhou (Institute of Biophysics, Chinese Academy of Sciences) for technical help with Biacore experiments.

