## Supplemental figures and extended methods for "Protective human anti-poxvirus monoclonal antibodies are generated from rare memory B cells isolated by multicolor antigen tetramers"

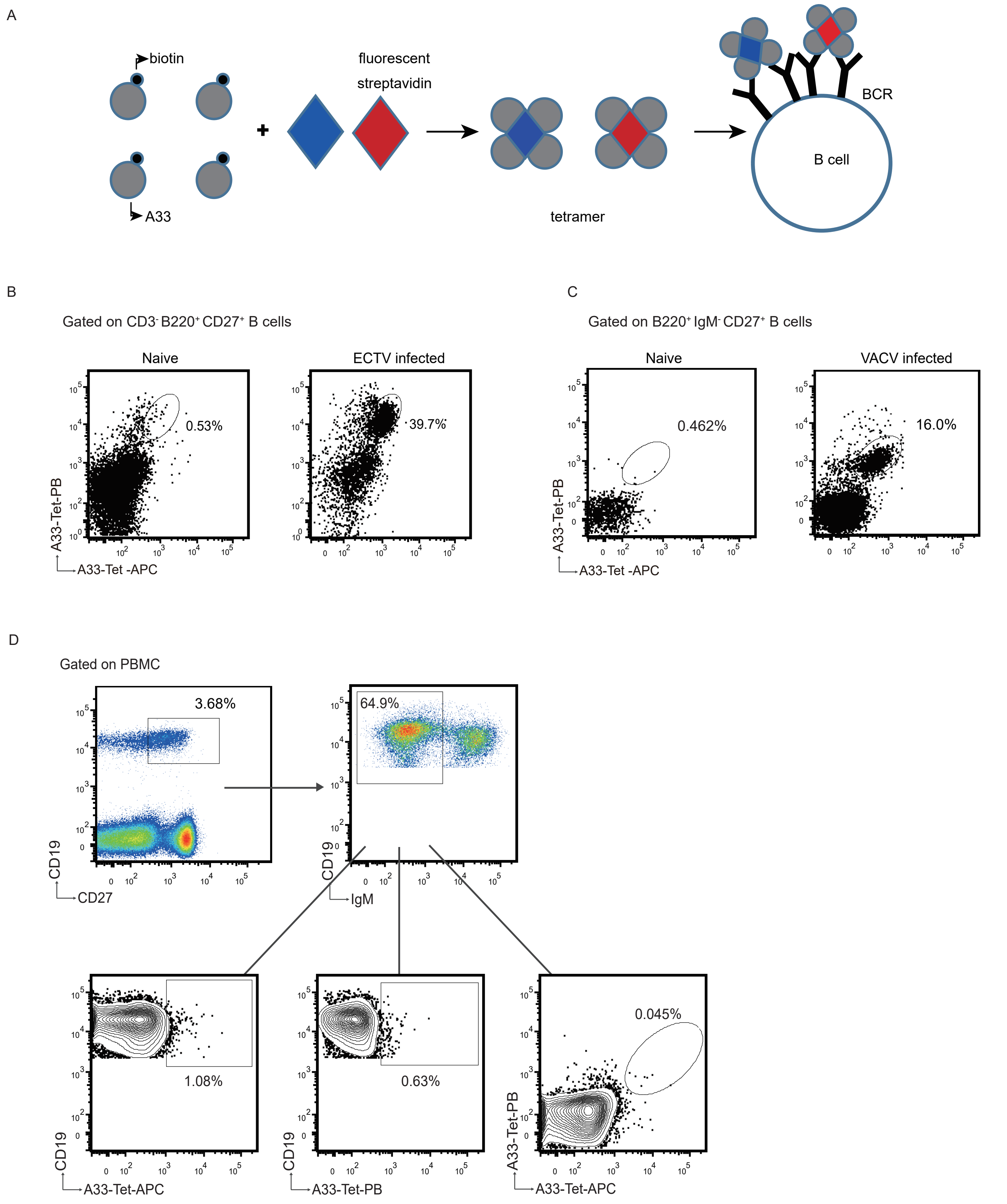


**Figure.S1 Generation of A33 tetramer and tetramer-based screening of antigen-specific B cells.** (A) Scheme of A33 tetramerizaion and B cell staining. Biotinylated A33 was incubated with different fluorescently labeled streptavidin to generate tetramers, respectively. Multiple A33 tetramers were used together along a panel of mAbs to indentify specific memory B cells. (B-C) Detection of A33-specific B cells in mice. (B) B6 mice were infected with 3000 PFU ECTV in the footpad. Splenocytes were isolated from ECTV infected mice at 4-5 weeks pi and stained for A33 specific B cells. Splenocytes from naive mice were used as control. The cells were first gated on $CD3^-$ $B220^+$ $CD27^+$ B cells, then A33-Tet-PB and A33-Tet-APC double positive B cells were identified. Numbers adjacent to gate represent the percentage of A33 tetramers positive cells within the gated B cells. (C) B6 mice were i.p. infected with $5\times10^6$ PFU VACV. Splenocytes were isolated from VACV infected mice at 4-5 weeks pi and stained for A33 specific B cells. Splenocytes from naive mice were used as control. The cells were first gated on $B220^+$ $IgM^-$ $CD27^+$ class switched memory B cells, then A33-Tet-PB and A33-Tet-APC double positive B cells were identified. Numbers adjacent to gate represent the percentage of A33 tetramers positive cells within the gated B cells. (D) Sorting of single A33-specific memory B cells from PBMC of a healthy volunteer. Identification of single A33 tetramer positive B cells or multicolor A33 tetramers positve B cells were shown in the gated $CD19^+$ $CD27^+$ $IgM^-$ class switched memory B cells. Frequency of each cell population is shown beside the population.

A

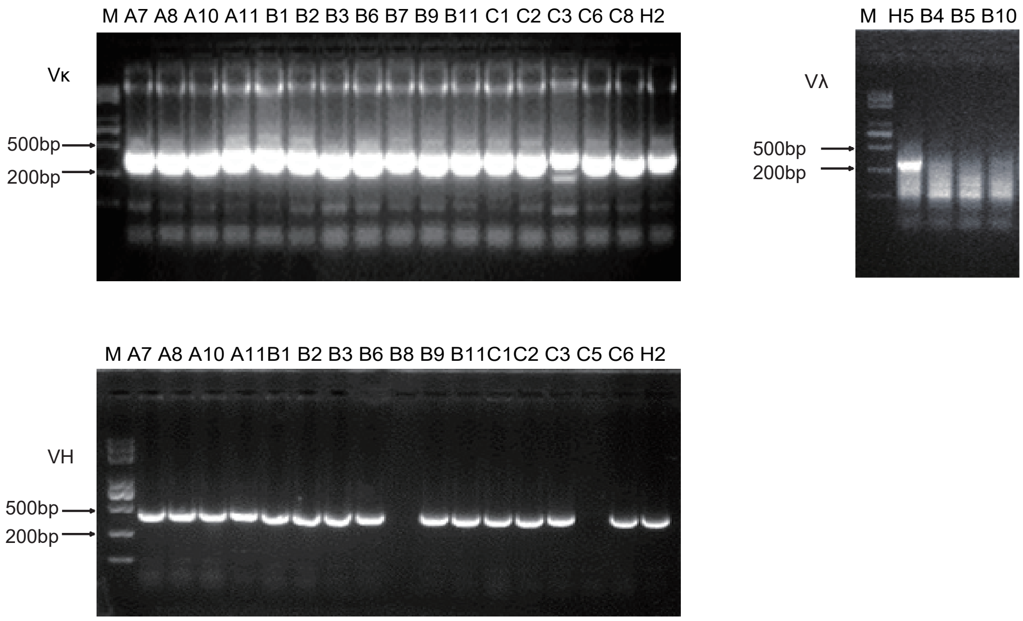


B

| | V-Gene and allele | | J-Gene and allele | |
|---|---|---|---|---|
| IgH | 14 | IGHV3-7*05 | 14 | IGHJ5*01 |
| | 4 | IGHV3-23*04 | 3 | IGHJ4*02 |
| | 1 | IGHV3-33*01 | 1 | IGHJ5*02 |
| | | | 1 | IGHJ3*02 |
| IgK | 11 | IGKV1-5*03 | 11 | IGKJ2*01 |
| | 8 | IGKV1-12*01 | 6 | IGKJ1*01 |
| | | | 2 | IGKJ5*01 |

C

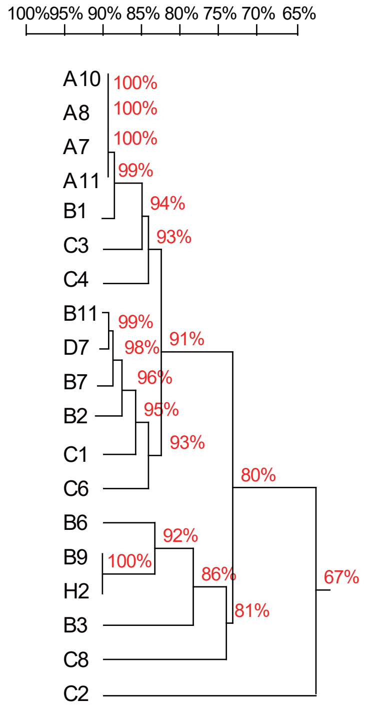


D

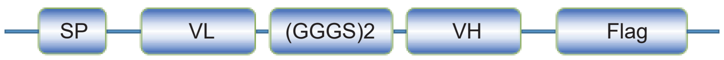


E

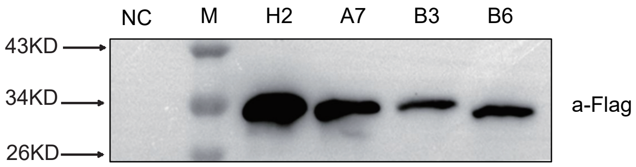


**Figure.S2 Amplification of Ab variable region and construction of ScFvs.** (A) VH and VL genes were amplified by nested RT-PCR from sorted single cell cDN Analyzed on a 2% agarose gel and cloned into pcDNA3.1 plasmid. (B) V, J regions of the 19 Abs were analyzed by IMGT. (C) DNAMAN software compared the sequence homology of paired VH and VL. (D) Scheme of consturct for scFv. (E) 293F cells were transfected with vector (negative control) or the indicated scFv plasmids. After 48 hours, the expression of scFvs was determined in the cell supernatant by western blot using the anti-flag antibody.

### Extended Materials and Methods

#### Cells, viruses and antibodies

293T cells (ATCC, CRL-3216) and BSC-1 cells (ATCC, 3168) were cultured in DMEM medium supplemented with 10% FBS, 2 mM L-glutamine, 100 μm nonessential amino acids, 10 mM HEPES buffer, 100 IU/ml penicillin, 100 μg/ml streptomycin (all from Gibco), and 0.05 mM β-mercaptoethanol (Amresco). 293-F cells (Invitrogen, R790-07) were cultured in SMM 293-TI (Sino-Biological). All cells were cultured at 37°C in 5% $CO_2$.

Initial stocks of ECTV Moscow and VACV Western Reserve were obtained from Dr. Luis Sigal (Thomas Jefferson University) and amplified and quantified as described (23). For the determination of virus titers in mice spleens, the spleens were made into a single cell suspension between two frosted slides and resuspended in 5 ml complete RPMI medium. 500μl of the cell suspensions were frozen and thawed three times and titers were determined in 10-fold serial dilutions of the cell lysates. Virus titers were calculated as PFU/spleen. To determine the virus titers in liver, a portion of the liver was weighed and homogenized in the medium using a tissue homogenizer and titrated.

Antibodies used for flow cytometry were anti-human CD19 (Biolegend,302208), anti-human IgM (Biolegend,31450), anti-mouse/rat/human CD27 (Biolegend,124214), anti-mouse CD8a (Biolegend，53-6.7), anti-mouse CD4 (Sungene, M10043-02E). anti-mouse IFN-γ (BD, 554412), anti-mouse B220 (Biolegend,103206), anti-mouse CD3 (Biolegend，100312), anti-mouse IgM (Biolegend，406507).

#### Production and Biotinylation of A33

The coding sequences for the extraviral domain of VACV Western Reserve A33 (24) were amplified by PCR from genomic DNA. For A33 the forward and reverse primers used were 5’AAACCATGGGCCATCACCATQCACCATCACTGCATGTCT-GCTAACGAGGCTG3’ and 5’AAAGGATCCGTTCATTGTTTTAACACAAAAATA CTTTC3’, respectively. To facilitate purification, the 6×His tag was fused to the

N-terminus. Avi-tag (GLNDIFEAQKIEWHE)(30) was fused to the C-terminus of A33 to mediate biotinylation. The whole construct was cloned into the PET-28a(+) vector and transformed into DH5α competent cells. The expression vector was verified by DNA sequencing and then transformed into BL21 (DE3) competent cells for expression. The transformed BL21 (DE3) cells were grown overnight at 37℃ and inoculated at 10% into LB medium. The culture was grown at 37℃ until the OD reached 0.6-0.8, IPTG (final concentration 0.4 mM) was added to induce protein expression and cells were harvested 4h later and lysed by sonication. The inclusion bodies were pelleted by centrifugation for 20 min at 8000×g, washed with washing buffer (20 mM Tris–HCl pH 8.0, containing 1% Triton-X100) followed by distilled water to remove contaminating salts and detergents.

Recombinant A33 was purified under denaturing conditions by Ni-NTA metal affinity chromatography. The inclusion bodies were solubilized with 8M urea and loaded onto Ni-NTA agarose (Thermo, QB211702), according to the manufacturer's recommendations. After washing out the unbound proteins, the target protein was eluted by 0.5M imidazole in 8M Urea lysis buffer. The purified proteins were refolded by dialysis against PBS (phosphate-buffered saline). Protein concentrations were determined using a bicinchoninic acid assay (EpiZyme, ZJ102) with bovine serum albumin as a standard. The purity of each protein was confirmed by sodium dodecyl sulfate-polyacrylamide gel electrophoresis (SDS-PAGE).

Biotinylation of purified A33 protein was perfomed by using the biotin-protein ligase kit (GeneCopoeia™, B1001) according to the manufacturer's insturction. Biotinylation of the protein wes verified by Western Blot using the HRP conjugated stretavidin (Earthox, E030100).

The coding sequences for the extraviral domain of VARV A33 with N terminus 6×His tag and C-terminus Avi-tag were synthesized by GenScript and directly cloned into the PET-28a(+). The purification and biotinylation of VARV A33 were the same as that of VACV A33.

**Flow cytometry and tetramer preparation**

To detect A33-specific B cells in mice, single-cell suspensions were prepared from spleens obtained from naive B6 mice, B6 mice infected with ECTV or B6 mice infected with VACV for 4-5 weeks. Red blood cells (RBC) were lysed with 0.84% $NH_4Cl$, the leukocytes were washed, and $2\times10^6$ leukocytes were stained with the indicated surface antibodies at 4°C for 30 min. Cells were analyzed with a BD LSRFortessa flow cytometer (BD Biosciences) and the data were analyzed using FlowJo (Tree Star, Ashland, OR).

All the tetramers were prepared freshly for each experiment. Biotinylated A33 was incubated with APC-conjugated streptavidin (SA-APC，eBioscience,17-4317-82), APC/Cy7-conjugated streptavidin (SA-APC/Cy7, Biolegend, 405208) or pacific blue-conjugated streptavidin (SA-PB, eBioscience，48-4317-82) for 3 min at room temperature at a molar ration of 4:1, respectively. The fluorescent A33 tetramers were then used together with other antibodies for cell staining.

To determinate the T cell responses, BALB/c mice were treated with H2 IgG or PBS before VACV infection. Seven days after the infection, mice were euthanized and single-cell suspensions of each spleen were prepared in 5ml complete RPMI medium. Following osmotic lysis of red blood cells with 0.84% $NH_4Cl$, the spleen cells were washed, and $10^6$ cells were stimulated for 6 h at 37℃ with $2\times10^5$ VACV infected or uninfected A20 cells in 96-well plates. Brefeldin A (BFA, Sigma) was added after 4.5 h to block the secretory pathway and allow for the accumulation of cytokines inside the cells. The cells were then stained for cell surface molecules, fixed, permeabilized, and stained for intracellular molecules using the Cytofix/Cytoperm kit (Becton Dickinson, BD) according to the manufacturer’s instructions. Cells were analyzed with a BD LSRFortessa flow cytometer and the data were analyzed using FlowJo.

**Single-cell sorting**

Blood samples were donated by a healthy volunteer with vaccination for smallpox at more than 40 years ago. Peripheral blood mononuclear cells (PBMC) were isolated from blood and then stained with the indicated cell surface markers. Memory B cells were gated as $CD19^+$ $IgM^-$ $CD27^+$. SA-APC $A33^+$ and SA-PB $A33^+$ double positive memory B cells were sorted as single cells into 96-well PCR plates containing 20μl/well of RT reaction buffer that included 5μl of 5× First strand cDNA buffer, 0.5μl of RNAseOut (Invitrogen, Carlsbad, CA), 1.25μl of DTT, 0.0625μl of Igepal and 13.25μl of dH2O (Invitrogen, Carlsbad, CA)(30). The plates were briefly centrifuged and immediately stored at -80°C until further processing. Cells were sorted on a BD FACS Aria IIIu and the data were analyzed using FlowJo.

**Constructing and expression of single chain antibody (scFv)**

The scFvs were constructed by connecting paired $V_L$ and $V_H$ with a $(GGGS)_2$ linker, an Ig leader sequence (METDTLLLWVLLLWVPGSTGD) was used as a signal peptidet at the N-terminus of scFv to mediate the secretion of the scFv into the cell supernatant. A flag tag was fused to the C-terminus of scFv for easy detection. The constructs were cloned into pcDNA3.1 vector (Invitrogen, V79020). The constructed scFvs were verified by sequencing. The plasmids were transfected into 293F cells with Polyethylenimine (PEI), and the cell supernatants were collected after 48-72 hours.

**Production of H2 IgG**

H2 IgG was generated by combining $V_H$ and $V_L$ with constant region of IgG1 heavy and light chain, respectively. The plasmids were synthesized by GenScript. The constructed heavy and light chain plasmids were co-transfected into 293F cells. The supernatant was collected and H2 IgG was purified by protein A (Nanomicro, UniMab 50) according to the manufacturer's recommendations. The purified protein was further dialysed with PBS. Protein concentration was determined using a bicinchoninic acid assay (EpiZyme, ZJ102) with bovine serum albumin as a standard. The protein was stored in aliquots at -80°C.

**ELISA**

96-well bottom ELISA plates were coated with VACV A33, VARV A33 (50 μg/ml), ECTV ($1\times10^7$ PFU/ml), VACV ($1\times10^7$ PFU/ml), PR8 ($1\times10^7$ PFU/ml) or BSA (50 μg/ml) at 4°C overnight as indicated. Plates were blocked for 2 h at 37°C with PBS containing 5% BSA. The plate were then incubated with the indicated antisera, cell culture supernatant or H2 IgG at 37°C for 2 hours. The plates were washed three times with PBS containing 0.05% Tween-20. For detection of sera, HRP-conjugated goat anti-mouse IgG (Jackson ImmunoResearch, 115-035-003) were added to each well and incubated for 1 h at 37°C. For detection of scFv, anti-flag M2 mAb (Sigma-Aldrich, F1804) were added to each well and incubated at 37°C for 2 h, and then washed five times. Then HRP-conjugated goat anti-mouse IgG (Jackson ImmunoResearch, 115-035-003) antibodies were added to each well and incubated for 1 h at 37°C. For detection of H2 IgG, HRP goat anti-human IgG (EARTH, E030170-02) were added to each well and incubated for 1 h at 37°C. After incubation with the HRP-conjugated antibodies, the plates were washed five times. TMB 1-Component Microwell Peroxidase Substrate (KPL) was added to each well and the plates were incubated at room temperature for 5-20 min. The reactions were stopped by adding 50μl 0.5 M $H_2SO_4$. The optical density (OD) at 450 nm was determined by Thermo MULTISKAN FC.

**Comet inhibition assay**

Confluent BSC-1 cells in 6 well plates were infected with ECTV (MOI=0.01) in 0.5 ml RPMI medium with 2.5% FBS (RPMI 2.5). The media containing virus was aspirated after 1h incubation at 37°C, and the cells were overlaid with 1.5 ml RPMI 2.5 containing the indicated dilutions of antisera or antibodies. Cells were incubated for 5 days at 37°C and stained with crystal violet as described for virus titers.

**Surface plasmon resonance analysis**

The affinity between H2 and A33 was measured at room temperature using a Biacore T100 system with CM5 chips (GE Healthcare). The immobilization of H2 was carried out following the canonical amino coupling chemistry, operating at a flow rate of 30 μL/min and injection for 60 seconds, dissociation for 180 seconds. A33 in varied concentrations (from 1μM to 62.5 nM, repeat at 250 nM) were allowed to flowover sensor surface in a running buffer of phosphate-buffered saline (PBS, pH 7.4) containing 0.01% (v/v) Tween-20. The binding affinity KD (equilibrium dissociation constant, KD = Kd/Ka, where Kd and Ka represent the dissociation rate constant and association rate constant, respectively) values were obtained using a series of A33 concentrations. The affinity was calculated using a 1:1 (Langmuir) binding fit model with BIA evaluation software.

**Statistics**

Statistical analysis was performed using Prism software (GraphPad). All statistical analyses were performed using an unpaired two-tailed Student's t-test or two-way ANOVA test as applicable. When applicable, data were displayed as mean ± SEM. Unless indicated, all displayed data correspond to one representative experiment of at least three similar independent experiments.
